# Production of extracellular polymeric substances in granular sludge under selection for *Accumulibacter* and *Competibacter*

**DOI:** 10.1101/2023.03.24.534144

**Authors:** Lorena B. Guimarães, Nina R. Gubser, Yuemei Lin, Jure Zlópasa, Simon Felz, Sergio Tomás Martínez, Mario Pronk, Thomas R. Neu, Morten K. D. Dueholm, Mads Albertsen, Rejane H. R. da Costa, Per Halkjær Nielsen, Mark C. M. van Loosdrecht, David G. Weissbrodt

**Affiliations:** Department of Biotechnology, Delft University of Technology, Delft, The Netherlands; Department of Chemistry and Bioscience, Center for Microbial Communities, Aalborg University, Aalborg, Denmark; Department of Sanitary and Environmental Engineering, Federal University of Santa Catarina, Brazil; Microbiology of Interfaces, Department River Ecology, Helmholtz Centre for Environmental Research – UFZ, Magdeburg, Germany; Department of Biotechnology and Food Science, Norwegian University of Science and Technology, Trondheim, Norway

**Keywords:** Exopolymers, Granular sludge, Microbiome, PAOs and GAOs, Wastewater treatment, Waste to product

## Abstract

Granular sludge intensifies the removal of nutrients from wastewater. Granules structured by extracellular polymeric substances (EPS) can be recovered as biomaterial. Links between microbial selection and EPS formation during granulation need to get uncovered. We inoculated anaerobic-aerobic sequencing batch reactors with either flocs or granules to study the relationships between microbial selection, bioaggregation, exopolymer formation, and EPS composition. Selection for slow-growing organisms like the model polyphosphate-accumulating organism “Candidatus Accumulibacter” (max. 83% vs. amplicon sequencing read counts) and glycogen-accumulating organism “Ca. Competibacter” (max. 45%) sustained granulation. Gel-forming exopolymers were produced as high as above 40% of the volatile solids of the biomass by stepwise increase of the organic loading rate (0.3 to 2.0 g COD_Ac_ d^-1^ L_R_^-1^). Confocal laser scanning microscopy, FT-IR spectroscopy, and HPAE-PAD chromatography revealed the complex and dynamic chemical compositions of the structural EPS in relation to microbial population shifts along reactor regimes. The analysis of 20 representative genomes of “Ca. Accumulibacter” and “Ca. Competibacter” recovered from public databases revealed their functional potential to produce EPS among other representative wastewater microorganisms. The more than 40 functional gene categories annotated highlight the complexity of EPS metabolic networks from monomers processing to assembly, export, and epimerizations. The combination of ecological engineering principles and systems microbiology will help unravel and direct the production of EPS from wastewater, valorizing residual granular sludge into beneficial biomaterials for the circular economy.

**Highlights:**

- Selection for slow-growing organisms like PAOs and GAOs fostered a robust granulation.
- Structural EPS were produced above 40% of biomass volatile content under high loading.
- Chemical composition of EPS evolved together with the microbial community composition.
- Genomic insights highlighted the genetic potential of PAOs and GAOs for EPS formation.
- Microbial communities are complex; further are their EPS compositions and metabolisms.

**Graphical abstract:** 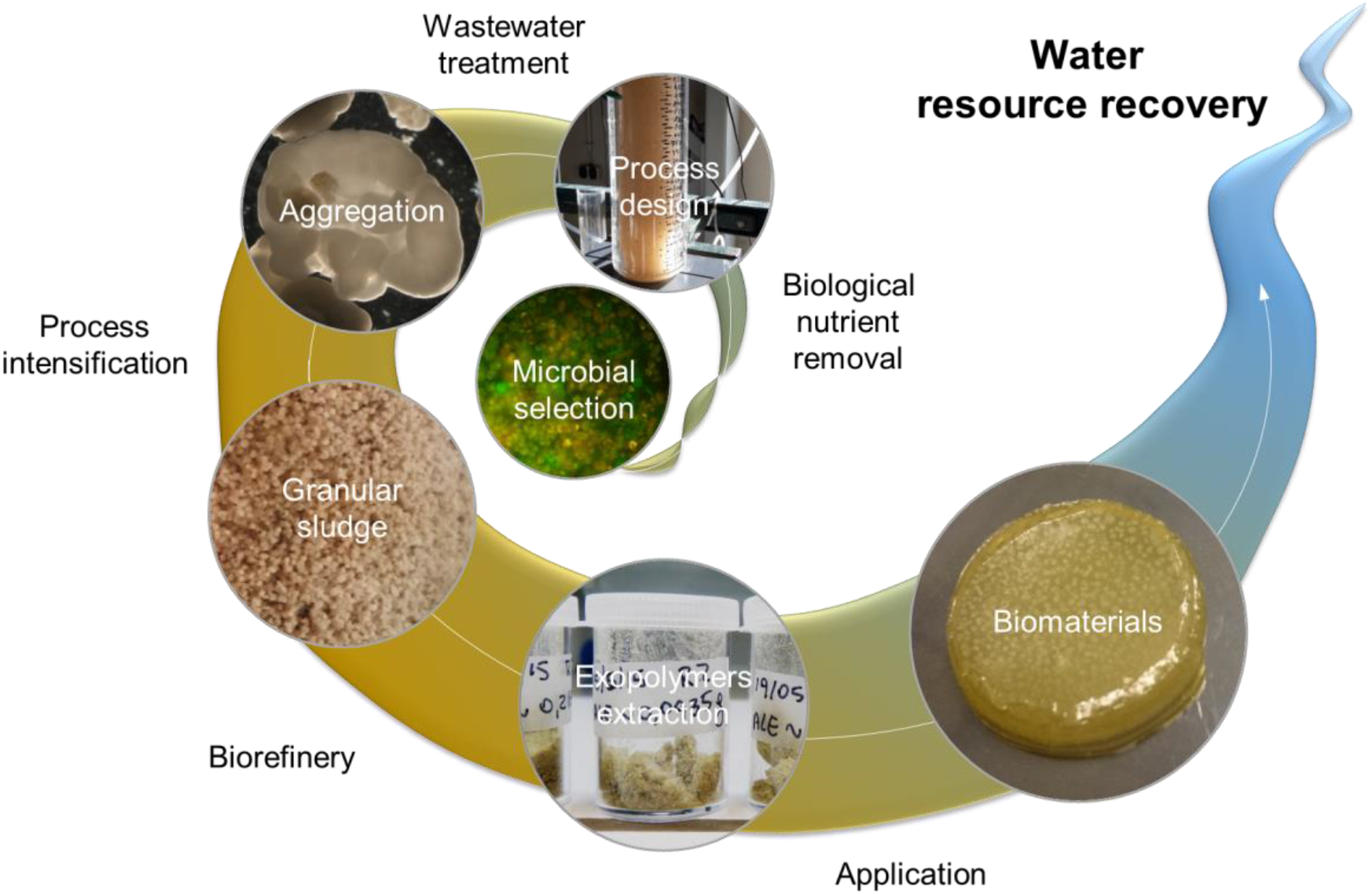

## 1 Introduction

The wastewater engineering sector faces new challenges for an integrated management of water, energy, and chemical resources in the circular economy (van Loosdrecht and Brdjanovic, 2014; Weissbrodt et al., 2020b). Granular sludge sequencing batch reactors (SBRs) (Giesen et al., 2012; Winkler and van Loosdrecht, 2022) made breakthrough to intensify the biological nutrient removal (BNR) from wastewater. These processes use mobile and fast-settling granular biofilms called “granules” formed by self-immobilization (Morgenroth et al., 1997) of activated sludge microorganisms (Ebrahimi et al., 2010; Weissbrodt et al., 2013a).

Granules support novel waste-to-resource approaches to recover extracellular polymeric substances (EPS) from the organic matter of wastewater (Lin et al., 2010). These exopolymers exhibit some similarities to bacterial alginates, like the complexing of metal ions, hydrogel formation, as well as hydrophilic and hydrophobic interactions. They can find hi-tech material applications, *e.g.*, in the coating, concrete, paper, nanocomposite, and flame-retardant industries (Lin et al., 2015; Zlopasa et al., 2015). Following the successful implementation of granular sludge for BNR in wastewater treatment plants (WWTPs) (Pronk et al., 2015b), a first process has been installed to produce exopolymers from waste granular sludge at full scale (Dutch Water Sector, 2019).

The formations of biofilms and exopolymers is widespread across microorganisms (Flemming and Wingender, 2010). Granules comprises a patchwork of EPS related to proteins, glycoproteins, amino sugars, and large glycoconjugates like nonulosonic acids, sialic acids, hyaluronic acids or sulfated glycosaminoglycans (Seviour et al., 2012; de Graaff et al., 2019; Boleij et al., 2020; Felz et al., 2020; Kleikamp et al., 2020). EPS extracted from granular sludge matrices and characterized using physical-chemical method, remain a challenge with respect to analytical biases (Felz et al., 2016; 2019; Seviour et al., 2019). Chemical compositions of EPS are complex, resulting from microbial and metabolic diversities (Weissbrodt et al., 2013a).

The compact structure of granules is stabilized by slow-growing microbial populations like polyphosphate (PAOs) and glycogen-accumulating organisms (GAOs) besides nitrifiers or anammox populations (van Loosdrecht et al., 1995; de Kreuk and van Loosdrecht, 2004). PAOs are selected for an enhanced biological phosphorus removal (EBPR) (Barnard and Abraham, 2006; Oehmen et al., 2010). Some PAO and GAO populations can denitrify when nitrate or nitrite are available as terminal electron acceptors (Rubio-Rincon et al., 2017). Because of their slower growth rate when compared to ordinary heterotrophic organisms (OHOs), PAOs and GAOs develop compact microcolonies (Larsen et al., 2006; Weissbrodt et al., 2013a). The PAO betaproteobacterial genus “*Candidatus* Accumulibacter” and the GAO gammaproteobacterial genus “*Ca.* Competibacter” (Hesselmann et al., 1999; Crocetti et al., 2002) are model populations in laboratory investigations of EBPR processes (Lopez-Vazquez et al., 2009; Weissbrodt et al., 2013b). Operation with real wastewater streams can select for other PAO and GAO populations depending on local wastewater compositions and environmental and operational constrains (Nielsen et al., 2012; Stokholm-Bjerregaard et al., 2017).

The diversity of microorganisms producing EPS in activated sludge and granular sludge remains unchartered. The physiology of PAOs and GAOs, and slow-growing organisms in general, and even more for exopolymer formation is unexplored. From the ecosystem model of BNR granular sludge (Weissbrodt et al., 2014b; Winkler et al., 2018; Weissbrodt et al., 2020a), “*Ca.* Accumulibacter” has been hypothesized to display metabolic traits for EPS biosynthesis, although often described as a slow-growing chemoorganoheterotroph. First sets of metabolic and enzymatic investigations of EPS formation in PAO enrichments have been recently reported (Tomás-Martínez et al., 2021; 2022; 2023a; 2023b). “*Ca.* Accumulibacter” belongs to the family of *Rhodocyclaceae* that comprises genera like *Zoogloea* and *Thauera* with well-known functionalities for EPS, biofilm and granule formations (Allen et al., 2004; Dugan et al., 2006; Weissbrodt et al., 2013a; Szabo et al., 2017; Guimarães et al., 2018). “*Ca.* Competibacter” may contribute to EPS formation in granules as well: the “granulan” polysaccharide has been characterized from a GAO-enriched granular sludge (Seviour et al., 2012), although accounting for only 2% of the mass of EPS extracted. Mass fractions of 0.15-0.30 g crude EPS g^-1^ dried biomass have been typically obtained from granules, while flocs harbor less than 0.10 g g^-1^ (Lin et al., 2013). PAOs and GAOs can sustain multiple objectives for process intensification and integration through granulation, BNR, phosphorus recovery (in case of PAOs), and EPS biorefinery (Stratful et al., 1999; Liao et al., 2003; Cornel and Schaum, 2009; Egle et al., 2016; Barnard et al., 2017).

We investigated the relationships between microbial selection, granulation, and EPS formation and compositions under conditions selecting for “*Ca.* Accumulibacter” and “*Ca.* Competibacter” in two replicated anaerobic-aerobic SBRs inoculated with either flocs or granules, respectively. Ecological engineering principles (Lawson et al., 2019) were applied to shape the microbiome of granular sludge and enhance the production of exopolymers. Mixed-culture biotechnology, structural microbiology, microbial ecology, chemical profiling of EPS, and analyses of EPS genetic signatures of publicly available (metagenome-assembled) genomes of PAOs, GAOs, and other representative lineages of wastewater environments were combined. We highlight that PAOs and GAOs are beneficial to both stabilize granular sludge and produce EPS at relatively high mass fractions, and further contribute to uncover the EPS biosynthetic potential of “*Ca.* Accumulibacter” and “*Ca.* Competibacter”.

## 2 Methods

### 2.1 Reactor operations for granulation and EPS formation under PAO/GAO selection

#### SBR operations

Two 2.4-L bubble-column SBRs of height-to-diameter ratios of 24 were inoculated in parallel at 1.5 g VSS L^-1^ with either activated sludge flocs (SBR1) or granular sludge (SBR2) collected from the BNR WWTPs Harnaschpolder and Garmerwolde (The Netherlands), respectively (**Fig. 1a**). Operational conditions for PAO-GAO selection were transferred from stirred-tank enrichment cultures (Weissbrodt et al., 2014a) into the granular sludge bubble-column reactor design. The SBRs were fed with acetate as sole carbon source and operated over 5 months at 18 ± 2°C and pH 7.5 ± 0.2 in cycles of 6 h under anaerobic-aerobic alternating conditions for a long-term evaluation of exopolymer formation and dynamics. The SBR cycles comprised a sequence of pre-feeding flush with dinitrogen gas (5 min), wastewater up-flow feeding (15 min), post-feeding flush with dinitrogen gas (5 min), anaerobic mixed batch (150-175 min), aerated mixed batch (150 min), biomass settling (30 to 5 min adapted as response to formation of a fast-settling biomass), and effluent withdrawal (5 min) at a volumetric exchange ratio of 50% (**Fig. 1b**).

**Figure 1.**
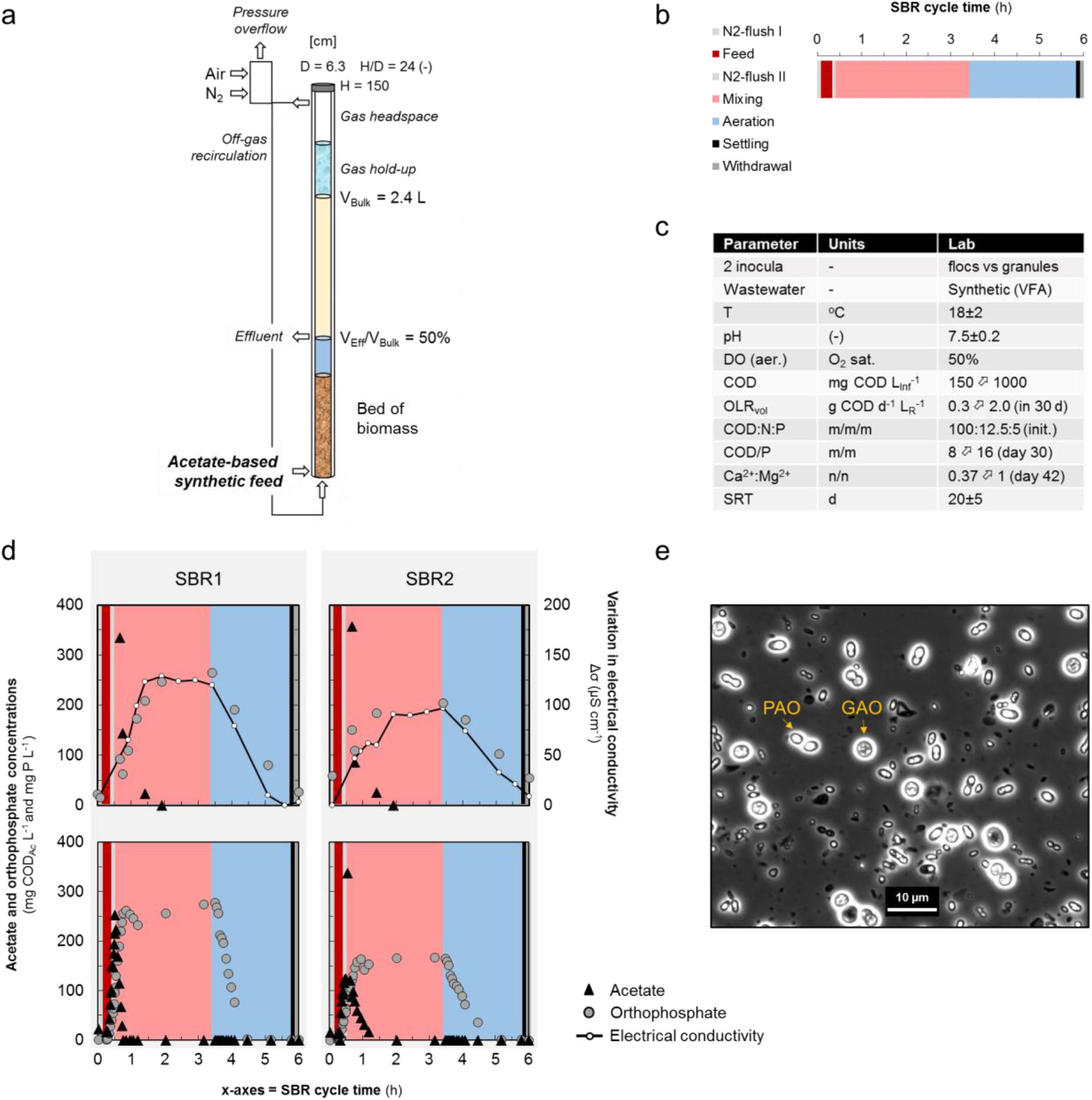
Design and operation of the SBRs. The bubble-column reactor design was equipped with off-gas recirculation loop and addition of air or dinitrogen to control the DO setpoints to 50% O2 sat. (**a**). The SBR cycle comprised an alternation of up-flow feeding of synthetic wastewater through the bed of granular sludge followed by an anaerobic batch phase and an aeration phase (**b**). SBR1 and SBR2 were seeded with flocculent activated sludge and granular sludge, respectively, from full-scale BNR installations and run at a SRT of 20 ± 5 days along with step-wise adaptation of the volumetric organic loading rate (OLR), COD/P ratio, and Ca^2+^:Mg^2+^ ratio (**c**). Overall, operational conditions were designed to foster a robust anaerobic selector for an efficient granulation under PAO/GAO selection by preventing leakage of COD into the aeration phase. This is exemplified for SBR1 and SBR2 under early operation (**d** – upper panels) and pseudo steady-state operation (**d** – lower panels). Electrical conductivity (absolute range of *ca.* 950-1250 μS cm^-1^ with the synthetic wastewater used) was powerful to monitor and manage the anaerobic selector and aeration phase length under the presence of PAOs. Electrical conductivity time course correlates directly with orthophosphate cycling (R^2^ = 0.90) and indirectly with acetate uptake. In lower d-panels, time series of acetate concentrations were measured at beds outlets during the 15 min of up-flow feeding of the wastewater across the beds (color allocation to cycle time periods is identical as in panel b). In these SBR cycle illustrations, the anaerobic mixing phase length can be decreased to < 2 h. Light microscopy image of PAOs and GAOs selected in SBR1 (**e**).

#### Feed characteristics

The up-flow feeding regimes of synthetic wastewater across the beds of biomass were achieved at a moderate flowrate of 0.080 L min^-1^, corresponding to a superficial flow velocity of 1.54 m h^-1^. The basic composition of the synthetic wastewater was identical to the one used for PAO enrichment by Weissbrodt et al. (2014a). The influent was supplemented with allyl-N-thiourea (ATU) to suppress nitrification and nitrite formation in order to prevent PAO inhibition. The start-up of SBR1 involved a step-wise increase of the volumetric organic loading rate (OLR) from 0.3 to 2.0 g COD_Ac_ d^-1^ L_R_^-1^ (*i.e.*, acetate converted as chemical oxygen demand – COD) in one month to guarantee a full anaerobic uptake of organics, while initially maintaining the nutrient ratios constant (8 g COD g^-1^ P and 20 g COD g^-1^ N-NH4). The Ca:Mg (0.37-1.00 mol Ca^2+^ mol^-1^ Mg^2+^), and C:P (8-16 g COD g^-1^ P) ratios were increased step-wise concomitantly to the relatively high final volumetric OLR in order to possibly increase the production of exopolymers. Similarly, the start-up of SBR2 involved adjustments of the volumetric OLR from 0.6-1.2 g COD d^-1^ L_R_^-1^ in 14 days and of the Ca:Mg (0.37-1.00 mol Ca^2+^ mol^-1^ Mg^2+^, day 50), and COD:P (8-16 g COD g^-1^ P, day 66) ratios (**Fig. 1c**).

#### Anaerobic selector characteristics

The anaerobic batch phase was designed as an anaerobic selector to ensure a complete uptake of organics prior to switching to aeration, in order to select for PAOs over OHOs (Weissbrodt et al., 2013a; Lochmatter and Holliger, 2014; Henriet et al., 2016). Mixing was achieved by up- flow recirculation of the reactor gas headspaces at 3.0 NL min^-1^ (superficial gas velocity of 0.016 m s^-1^) at the reactors feet using a gas pump N811 KN.18 (KNF Verder, Germany). Dinitrogen gas was injected at 1.0 NL min^-1^ in the headspace recirculation loops using mass flow controllers 5850TR series (Brooks, USA) in order to maintain anaerobic conditions in the bulk liquid phases. On-line electrical conductivity sensors (Metrohm, Switzerland) were used to monitor and control the length of the anaerobic phases as function of orthophosphate release from the anaerobic metabolic activity of PAOs (Aguado et al., 2006; Weissbrodt et al., 2014a). Residual concentrations of VFAs by the end of the anaerobic batch phases were measured (see below) and the anaerobic phase lengths adapted manually to prevent any leakage from the anaerobic into the aeration phase (**Fig. 1d**).

#### Aeration phase characteristics

The dissolved oxygen (DO) concentrations were measured with AppliSens low-drift sensors (Applikon Biotechnology, Netherlands) and controlled at 50% O2 saturation to maintain stable oxygenation conditions during aeration by injection of air or dinitrogen gas in the headspace recirculation loops. The aeration phase lengths were adjusted based on an on-line electrical conductivity measurement that correlated linearly with orthophosphate uptake by PAOs. The sludge retention time was maintained at 20±5 days by sampling and periodical manual purge of excess biomass from the aerated mixed liquors by the end of SBR cycles.

#### Settling characteristics

Granulation was driven by stimulating a spontaneous bioaggregation resulting from the selection of PAOs and GAOs that form compact microcolonies, similar to PAO/GAO enrichment cultures (Weissbrodt et al., 2013a). The settling time was adjusted in response to the fast-settling sludge that progressively formed and accumulated as a result of PAO and GAO selection, similar to earlier reports (Lochmatter and Holliger, 2014; Weissbrodt et al., 2014a), in order to minimize the traditional harsh wash-out conditions used for granulation at bench scale (Beun et al., 1999).

### 2.2 Analyses of global parameters

#### Measurements of dissolved components

Concentrations of acetate were measured from the bulk liquid phases using a high-performance liquid chromatograph (HPLC) equipped with UV and RI detectors (BioRad, USA), after sample filtration on 0.2 μm filters. Ammonium (NH4^+^-N), orthophosphate (PO4^3-^-P) and total nitrogen (TN) were measured spectrophotometrically using colorimetric kits (Hach-Lange, Germany).

#### Measurements of biomass

Concentrations of total (TSS), inorganic (ISS), and volatile (VSS) suspended solids were analyzed according to the Standard Methods (APHA-AWWA-WEF, 2012). Heights of biomass beds in the reactors were measured daily using tailor’s tape measures to manage the anaerobic contact time (Weissbrodt et al., 2017). Evolutions of particle size distributions in the biomass were followed using a Quantiment 500 Image Analyzer (Leica Cambridge Instruments, UK).

### 2.3 Extraction and mass fractions of exopolymers from granular sludges

#### Extractions of EPS

Exopolymers were extracted from 1 g of centrifuged and lyophilized mixed liquors that were sampled by the end of aeration phases. Extractions were achieved under conditions combining 30 min stirring (400 rpm), alkaline conditions (pH∼10) by addition of 0.5% (w/v) Na2CO3, and high temperature (80°C) for complete dissolution of granular biofilm matrices (Lin et al., 2010; Felz et al., 2016). Exopolymers were then precipitated under acidic conditions by stepwise addition of HCl at 1 mol L^-1^, at room temperature (23±2 °C), freeze-dried, and weighed.

#### Measurements of mass fractions of EPS

The mass fractions of the raw exopolymer extracts were measured as ratio of total solids per unit of biomass measured as volatile solids (TSEPS VS_BM_^-1^) over about 40 time points along the experimental runs. Inorganic and volatile fractions of 8 selected exopolymers extracts were analyzed together with their physical-chemical properties by thermogravimetry coupled to differential scanning calorimetry (TG/DSC) (Ukrainczyk et al., 2013). This led to the expression of volatile mass fractions of exopolymers in the biomass (VS_EPS_ VS_BM_^-1^). The TSEPS VS_BM_^-1^ ratio was considered to take into account the presence of minerals that stabilize the EPS matrix, while the VSEPS VS_BM_^-1^ ratio targeted the organic content of EPS.

### 2.4 Chemical and material characterizations of EPS

#### Mapping of predominant glycoconjugates in granule cross-sections

Glycoconjugates were mapped within granule cryosections by fluorescence lectin-binding analysis (FLBA) following Weissbrodt et al. (2013a) and de Graaff et al. (2019) via screening of green fluorescent lectins (Neu and Kuhlicke, 2017; 2022).

#### Gelling property of exopolymers

The gelling property of the exopolymer extracts was verified by formation of sodium-based thin films and calcium-based beads (Seviour et al., 2009; Lin et al., 2013).

#### Overall chemical compositions of EPS

Variations in overall chemical compositions of the extracted EPS were measured by Raman and FT-IR spectroscopy methods (Lin et al., 2010).

#### Analysis of protein and polysaccharide contents of EPS

The protein (PN) and polysaccharide (PS) contents of the lyophilized exopolymer extracts were analyzed by bicinchonic acid (BCA) and Dubois assays, respectively (Guimarães et al., 2018; Felz et al., 2019). Adaptations were made to the protocol on the Dubois method: the single glucose standard used for the colorimetric and spectrophotometric analysis was replaced by a mixture of sugar standards similar to the ones used for the high-resolution analysis by HPAE-PAD chromatography described hereafter. This helped to account for the variability in detection sensitivity of the multiple types of monomers present in the EPS. The PN and PS contents were used to calculate the protein-to-polysugar (PN/PS) ratio of the exopolymers.

#### Profiling of monosaccharides in the EPS

The monosaccharide analysis in the extracted exopolymers was performed according to Felz et al. (2019). Exopolymer samples were hydrolyzed for 8 h at 105°C with 1 mol L^-1^ HCl, at a concentration of 10 mg mL^-1^. Following the hydrolysis, samples were centrifuged at 10,000 × g for 5 min. The supernatant was collected, neutralized with 1 mol L^-1^ NaOH and diluted 1:5 with ultrapure water. The diluted samples were filtered through a 0.45 µm PVDF filter. The analysis of the hydrolyzed samples was performed by high-performance anion-exchange chromatography with pulsed amperometric detection (HPAEC-PAD) on a Dionex ICS 5000^+^ ion chromatograph equipped with an AminoTrap pre-column and a PA20 column (Thermo Fischer Scientific Dionex, USA). Standards of glycerol, fucose, galactosamine, rhamnose, glucosamine, galactose, glucose, xylose, mannose, ribose, galacturonic acid and glucuronic acid were spiked to identify the predominant compounds in the chromatograms.

### 2.5 Molecular analyses of bacterial community compositions and population dynamics

#### V1-V3 16S rRNA gene amplicon sequencing

Bacterial selection was measured at high throughput and resolution by analyzing time series of bacterial community compositions using V1-V3 16S rRNA gene-based amplicon sequencing, following the MIDAS field guide of activated sludge (McIlroy et al., 2015b). In short, genomic DNA was extracted from biomass samples collected every 2-3 days along the operation of SBR1 and SBR2 (*i.e.*, *ca.* 60-75 samples per reactor), using FastDNA SPIN Kits for Soil (MP Biomedicals, USA). The pair of forward 27F (5’-AGAGTTTGATCCTGGCTCAG-3’) and reverse 534R (5’-ATTACCGCGGCTGCTGG-3’) eubacterial primers were used to amplify the V1-V3 hypervariable region of the 16S rRNA gene by polymerase chain reaction (PCR). After purification on Agencourt AMPureXP beads (Becan Coulter, UK), the amplicons from the set of samples were pooled and sequenced at an average depth of 23639 ± 6476 reads per sample using a MiSeq desktop sequencer (Illumina, USA). The sequencing datasets were processed following the bioinformatics procedure summarized by Karst et al. (2016) and using the ampvis R package (Albertsen et al., 2015). Sequencing reads forming operational taxonomic units (OTUs) were mapped against the MiDAS database, and relative abundances of closest bacterial relatives were expressed as percentage of total sequencing read counts.

#### 16S rRNA targeted fluorescence in situ hybridization

Analyses of 16S rRNA-targeted fluorescence *in situ* hybridization (FISH) and epifluorescence microscopy (EFM) were conducted following Nielsen et al. (2016) to qualitatively validate shifts in predominant microbial populations. In short, mixed liquor samples were homogenized using a Potter-Elvejem tissue grinder. Cells were fixed, immobilized, and hybridized according to standard practice (Amann et al., 1995; Nielsen et al., 2009). Eubacteria were targeted by the EUB338mix set of oligonucleotides (Amann et al., 1990; Daims et al., 1999). “*Ca.* Accumulibacter” was first targeted using the PAOmix set of probes PAO462, PAO651, and PAO846 (Crocetti et al., 2000). The detection of PAOs was refined by only using the PAO651 probe, since other PAO462 and PAO846 hybridize to other closely related lineages (Albertsen et al., 2016). The PAO clades I and II were targeted by the probes Acc-1-444 and Acc-2-444, respectively (Flowers et al., 2009; Welles et al., 2015). “*Ca.* Competibacter” was targeted using the GAOmix set of probes GAO_Q431 and GAO_Q989 (Crocetti et al., 2002). Microscopy slides were analyzed using an Axioplan 2 epifluorescence microscope (Zeiss, Germany) equipped with filter sets 26, 20, and 17 for Cy5, Cy3 and FLUOS chromophores, respectively.

### 2.6 Genomic analysis of the EPS biosynthetic potential of “*Ca*. Accumulibacter” and “*Ca*. Competibacter” among other microbial populations of granular sludge

A set of representative genomes of “*Ca.* Accumulibacter”, “*Ca.* Competibacter”, and other microbial populations of granular sludge, as well as model organisms of exopolysaccharide production listed hereafter, were imported from public databases into KBase (Arkin et al., 2018), functionally annotated using the RAST toolkit (Brettin et al., 2015), and functionally profiled using the Domain Profile app. The functional genetic outputs were processed as a heatmap to display the genetic potential and (dis)similarities in EPS metabolism of the lineages.

#### Representative genomes of “Ca. Accumulibacter” and “Ca. Competibacter”

A representative set of 19 genomes of “*Ca.* Accumulibacter” was selected out of the more than 30 metagenome-assembled genomes (MAGs) deposited in public repositories which were recovered from a previous study (Rubio-Rincon et al., 2019) and analyzed in KBase (Allen et al., 2017). These representative genomes were selected based on the computation of whole-genome similarity metrics by average nucleotide identity (ANI) scores using the FastANI algorithm (considering that genome compositions are different when ANI < 95%) (Jain et al., 2018). Two additional MAGs from the GAO lineages “*Ca.* Competibacter phosphatis” and “*Ca.* Competibacter denitrificans” (McIlroy et al.) were added to the genome set.

#### Representative genomes of other microbial populations of granular sludge

Genomes from granular sludge microorganisms were imported to compare the genetic signatures of “*Ca.* Accumulibacter” and “*Ca.* Competibacter” into the broader context of the microbial ecosystem of BNR granular sludge (Weissbrodt et al., 2014b; Winkler et al., 2018; Weissbrodt et al., 2020a). This ecosystem model and an additional microbial ecology study of granules (Guimarães et al., 2018) were used to retrieve about 50 genomes of (*i*) other PAO genera like *Tetrasphaera* and “*Ca.* Halomonas”; (*ii*) other GAO genera like “*Ca.* Defluviicoccus seviourii”, “*Ca.* Contendobacter odensis” and *Thermomonas*; (*iii*) denitrifiers like *Zoogloea*, *Thauera*, *Dechloromonas*, *Methyloversatilis*, *Azoarcus* and *Azonexus* in the family of *Rhodocyclaceae* of “*Ca.* Accumulibacter; (*iv*) nitrifiers like *Nitrosomonas*, *Nitrospira*, *Nitrococcus*; as well as (*v*) other flanking populations present in BNR granular sludge, namely *Acidovorax*, *Acinetobacter*, *Aminobacter*, *Anabaena*, *Anaerolineae*, *Aquamicrobium*, *Aquimonas*, *Arcobacter*, *Bradyrhizobium*, “*Ca.* Brocadia”, *Chelatococcus*, *Chlorobiaceae*, *Chryseobacterium*, *Cloacibacterium*, *Clostridium*, *Comamonas*, *Cytophaga*, *Delftia*, *Devosia*, *Flavobacterium*, *Hydrogenophaga*, *Leptothrix*, *Lysobacter*, *Macellibacteroides*, *Microvirgula*, *Nitrococcus*, *Pirellula*, *Planctomyces*, *Pseudoxanthomonas*, *Rhodobacter*, “*Ca*. Saccharimonas”, *Simplicispira*, *Sphaerotilus*, *Stenotrophomonas*, *Sulfurospirillum*, *Thiothrix*.

#### Representative genomes of model organisms producing exopolysaccharides

The genomes of model organisms of exopolysaccharide production were used for comparison of sequence homologies within the genera *Anabaena*, *Azotobacter*, *Escherichia*, *Komagateibacter*, *Pseudomonas*, *Sphingomonas*, *Sinorhizobium*, *Streptococcus* and *Xanthomonas*, complementing Schmid et al. (2015).

## 3 Results

### 3.1 Granulation and EPS production was efficient under selection for PAOs and GAOs

#### Performance of the first SBR inoculated with flocculent activated sludge

In SBR1, inoculated with BNR activated sludge, granulation was achieved in the first month of anaerobic-aerobic operation on the acetate-based synthetic wastewater (**Fig. 1**), while minimizing biomass wash-out (**Fig. 2a**). A dense granulation was promoted by selection for slow-growing PAOs. The granular biomass exhibited a median particle diameter of 195 µm (1^st^-3^rd^ quartiles = 129-285 µm). The total suspended solids (TSS) accumulated from 1.5 to 6 g TSS L_R_^-1^. The content of volatile suspended solids (VSS) decreased from 75% to 57% of the TSS. The increase in inorganic suspended solids (ISS) to up to 43%, relating to the progressive selection for PAOs that accumulate inorganic polyphosphate. The mass fraction of exopolymers raised from 0.20 to 0.30 g TS_EPS_ g^-1^ VS_BM_ (that takes into account minerals that stabilize the EPS), or 0.14 to 0.21 g VS_EPS_ g^-1^ VS_BM_ in terms of volatile content of EPS (*i.e.*, only displaying the organic content of EPS in the biomass) (**Fig. 2b**). Under this initial period, GAOs transiently dominated (**Fig. 2c**) with “*Ca.* Competibacter”, “*Ca.* Contendobacter”, and CPB_S60 lineages making 3-23% of the total read counts in the V1-V3 16S rRNA gene amplicon sequencing dataset (**Fig. 3**), before PAOs became predominant.

**Figure 2.**
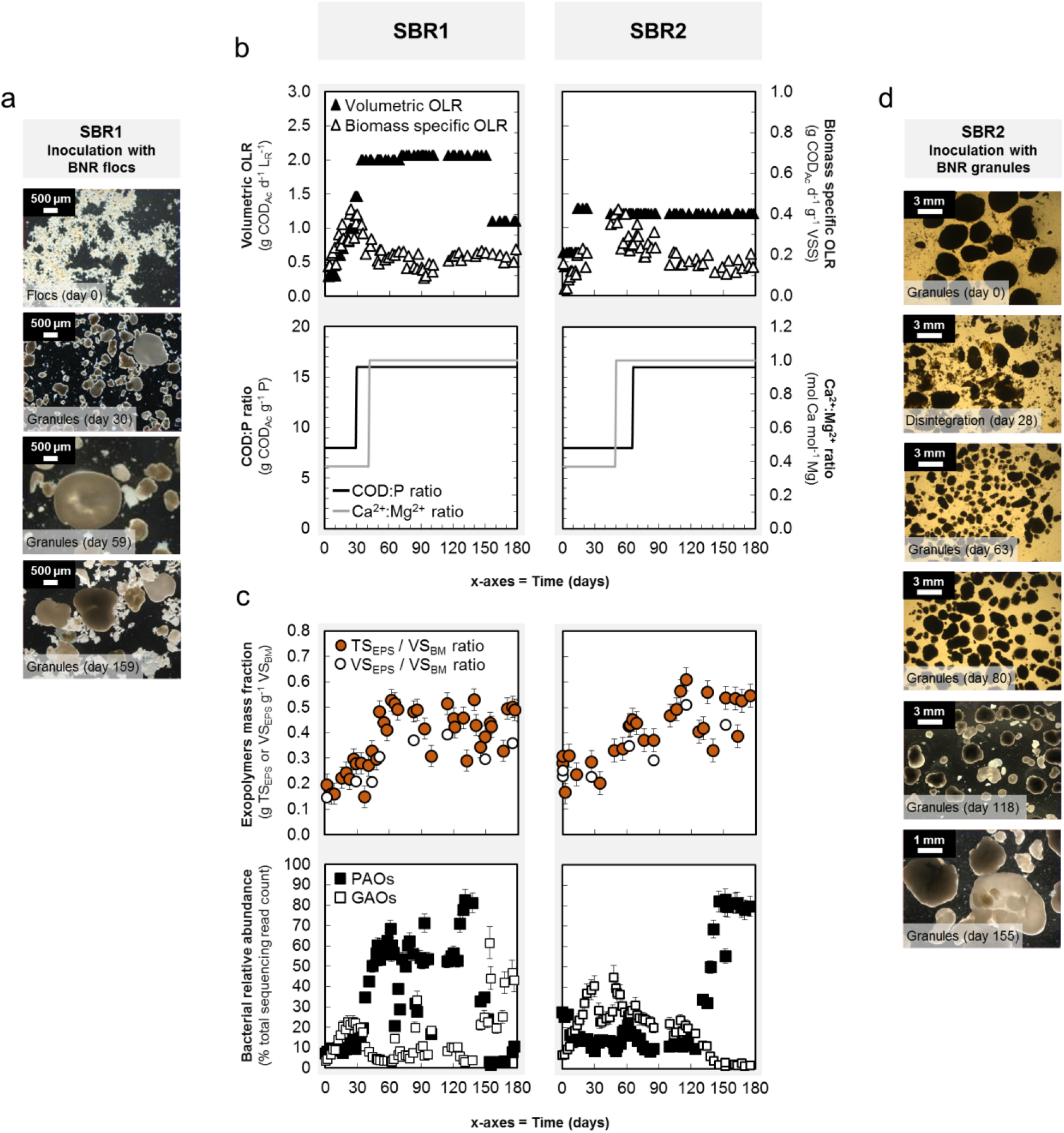
Granulation and exopolymer production. Granulation was achieved spontaneously by applying conditions selecting for PAOs and GAOs in SBR1 that was inoculated with activated sludge flocs (**a**). Operational conditions involved a stepwise increase in the volumetric organic loading rate (OLR) (*black triangles*) to ensure a full uptake of COD in the anaerobic selector, while the biomass specific OLR (*white triangles*) evolved freely in function of the biomass accumulation in the system (**b** – upper panels). The COD/P ratio (*black lines*) and Ca^2+^:Mg^2+^ ratio (*grey lines*) were over the first 2 months in order to sustain a good granulation (**b** – lower panels). Exopolymers were produced at substantial mass fractions reaching plateaus as high as 0.50 ± 0.1 g TS_EPS_ g^-1^ VS_BM_ (*orange dots*) or 0.40 ± 0.01 g VS_EPS_ g^-1^ VS_BM_ (*white dots*) (**c** – upper panels). GAOs (*white squares*) initially popped up in the two systems: they formed up to 20 % of the bacterial community over the first month of operation in SBR1 seeded with flocs, while remained predominant up to 40 % over almost 4 months in SBR2 seeded with granules; PAOs (*black squares*) then predominated the granular sludges up to 83 % of their bacterial communities (**c** – lower panels). After 5 months of operation of SBR1, a pH control issue disturbed the process and resulted in an almost half bed loss, requiring adaptation of a 2-fold lower volumetric OLR to maintain a stable biomass specific OLR. This led to a shift in predominance from PAOs to GAOs, but at no impact on the exopolymer mass fraction of the remaining granular sludge, highlighting the different response times of microbial populations (fast dynamics) *vs*. EPS matrices (resistance). Inoculation with full-scale granules in SBR2 resulted first in transient disintegration prior to regranulation (**d**).

**Figure 3.**
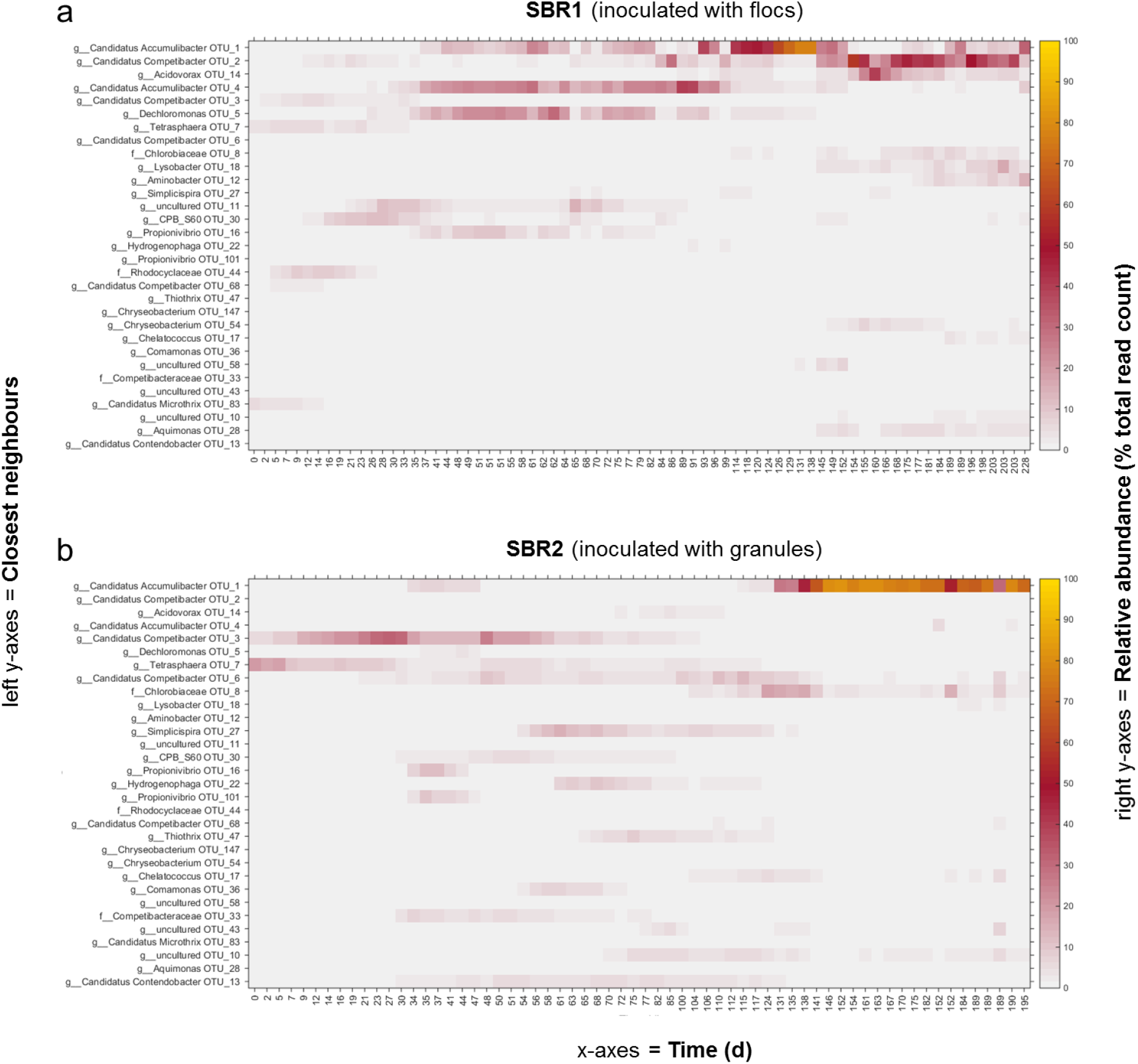
Heatmaps highlighting the dynamics of the predominant bacterial lineages (>5% total read count) such as the PAO “*Ca.* Accumulibacter” and GAO “*Ca.* Competibacter” measured from the microbiomes of the granular sludge biomasses of SBR1 after inoculation with BNR flocculent sludge dominated by *Tetasphaera* and “*Ca.* Microthrix” (**a**) and SBR2 after inoculation with BNR granular sludge dominated by *Tetrasphaera* and “*Ca.* Competibacter” (**b**) by V1-V3 16S rRNA gene-based amplicon sequencing. The predominant operational taxonomic units (OTUs; at least one instance > 5% in one of the two SBRs) were sorted in decreasing order of relative abundances across the two systems.

Exopolymers were significantly produced up to 0.48 g TS_EPS_ g^-1^ VS_BM_ (0.31 as VS ratio) during the second month along with the proliferation of PAOs from 12 to 68 %, after reaching the nominal volumetric organic loading rate (OLR) of 2 g COD_Ac_ d^-1^ L_R_^-1^ and after adjusting the Ca:Mg ratio from 0.37 to 1.00 mol Ca^2+^ mol^-1^ Mg^2+^ and the COD:P ratio from 8 to 16 g COD g^-1^ P. Meanwhile, granules exhibited a median particle size of 307 µm (483-781 µm). The biomass specific OLR decreased from 0.40 to 0.18 g COD_Ac_ d^-1^ g^-1^ VSS, as a result of biomass accumulation up to 8 g TSS L_R_^-1^ under constant nominal volumetric OLR. The main PAOs affiliated to the genus “*Ca.* Accumulibacter” accompanied by *Dechloromonas*, whereas *Tetrasphaera* declined after inoculation. In both SBRs, measurements by fluorescence *in* situ hybridization (FISH) qualitatively confirmed the decline of *Tetrasphaera* from the inoculation sludges to the transient predominance of GAOs and the enrichment in the PAO “*Ca*.

Accumulibacter” under pseudo steady-state conditions (not shown). Over the first 60 days, granulation was linearly correlated (r = 0.90) to the preferential enrichment of “*Ca.* Accumulibacter” and the exopolymer production.

A high mass fraction of exopolymers was successfully maintained at 0.46 g (0.41-0.49 g) TS_EPS_ g^-1^ VS_BM_ and 0.36 g/g (0.31-0.37 g/g) as VS ratio for more than 100 additional days under pseudo steady state with a stable bed of granular sludge dominated by “*Ca.* Accumulibacter” at 56% (52-60%; max. 83%). This highlighted the potential for production of exopolymers on the long run. The exopolymer matrix formed under such conditions was resistant, even in the course of a drastic disturbance in pH that impacted the reactor operation after 150 days: this resulted in a significant shift in predominance from PAOs to GAOs (up to 50-60%) while the exopolymer mass fraction remained constantly high, highlighting the product robustness.

The bacterial community (**Fig. 3**) further harbored major populations that accompanied the genus “*Ca*. Accumulibacter” during granulation and exopolymer production, namely the putative PAO *Dechloromonas* (0.1-22%) and putative GAO *Propionivibrio* (n.d.-10%) both affiliating with the family of *Rhodocyclaceae*, and *Flavobacterium* (0.3-8%). Populations of *Tetrasphaera* (8-0.1%), *Rhodobacter* (6-0.2%), and the filamentous “*Ca.* Microthrix” (11-0.01%) were predominant in the inoculum prior to declining during operation with the synthetic wastewater supplied with volatile fatty acids (VFAs). “*Ca.* Competibacter” (2-7%) and the candidate lineage CPB_S60 (0.2-10%) in the reappraised family of *Competibacteraceae* (McIlroy et al., 2015a) displayed a transient early selection during the first two weeks before declining along with the enrichment of “*Ca.* Accumulibacter”.

#### Performance of the second SBR inoculated with granular sludge

In SBR2, the granular sludge seeded from a full-scale installation disintegrated into small aggregates of 47 µm (37-63 µm) that re-granulated into particles of 107 µm (87-226 µm) in 2 months. GAOs dominated first (6-45%) with “*Ca.* Competibacter”, “*Ca*. Contendobacter” and CPB_S60. Exopolymers made 0.27 ± 0.06 g TS_EPS_ g^-1^ VS_BM_ (**Fig. 2b-c**, **Fig. 3**). The population of the genus *Simplicispira,* that may contribute to EBPR activity (Kleemann, 2015), raised from 1 to 13%, while GAOs were outcompeted over the next 2 months. The exopolymer mass fraction increased from 0.34 to as high as 0.61 g TS_EPS_ g^-1^ VS_BM_. *Propionivibrio* maximally reached 12% on day 35 before declining to undetectable levels. PAOs dominated since then from 30% to 83% in the amplicon sequencing dataset with “*Ca*. Accumulibacter” and *Dechloromonas* succeeding to *Tetrasphaera* which decreased from 27% to as low as 0.15%.

A sustainably high mass fraction of exopolymers (0.50 ± 0.1 g TS_EPS_ g^-1^ VS_BM_ and 0.40 ± 0.01 g g^-1^ as VS ratio) was obtained together with an optimum formation of granule particles of 768 µm (438-1170 µm). Similar to SBR1, the stepwise increase in the volumetric OLR up to 1.2 g COD_Ac_ d^-1^ L_R_^-1^ in 15 days and the adjustments made in the Ca:Mg ratio from 0.37 to 1.00 mol Ca^2+^ mol^-1^ Mg^2+^and in the COD:P ratio from 8 to 16 g COD g^-1^ P, sustained the granulation and exopolymer formation. The biomass content slowly increased up to 8 g VSS L^-1^ in 150 days, with an ash content stabilizing at 45 ± 4%. The biomass specific OLR increased up to 0.42 g COD_Ac_ d^-1^ g^-1^ VSS in 2 months, prior to decreasing over the next 60 days and maintaining at 0.19 (0.15-0.23) g COD_Ac_ d^-1^ g^-1^ VSS.

The characteristics of granules, exopolymer mass fractions, and predominant populations obtained under operation of SBR1 and SBR2 are summarized in **Table 1**. Overall, a high production of exopolymers was achieved in both SBRs together with a robust granulation state under selection for PAOs and GAOs.

**Table 1.**
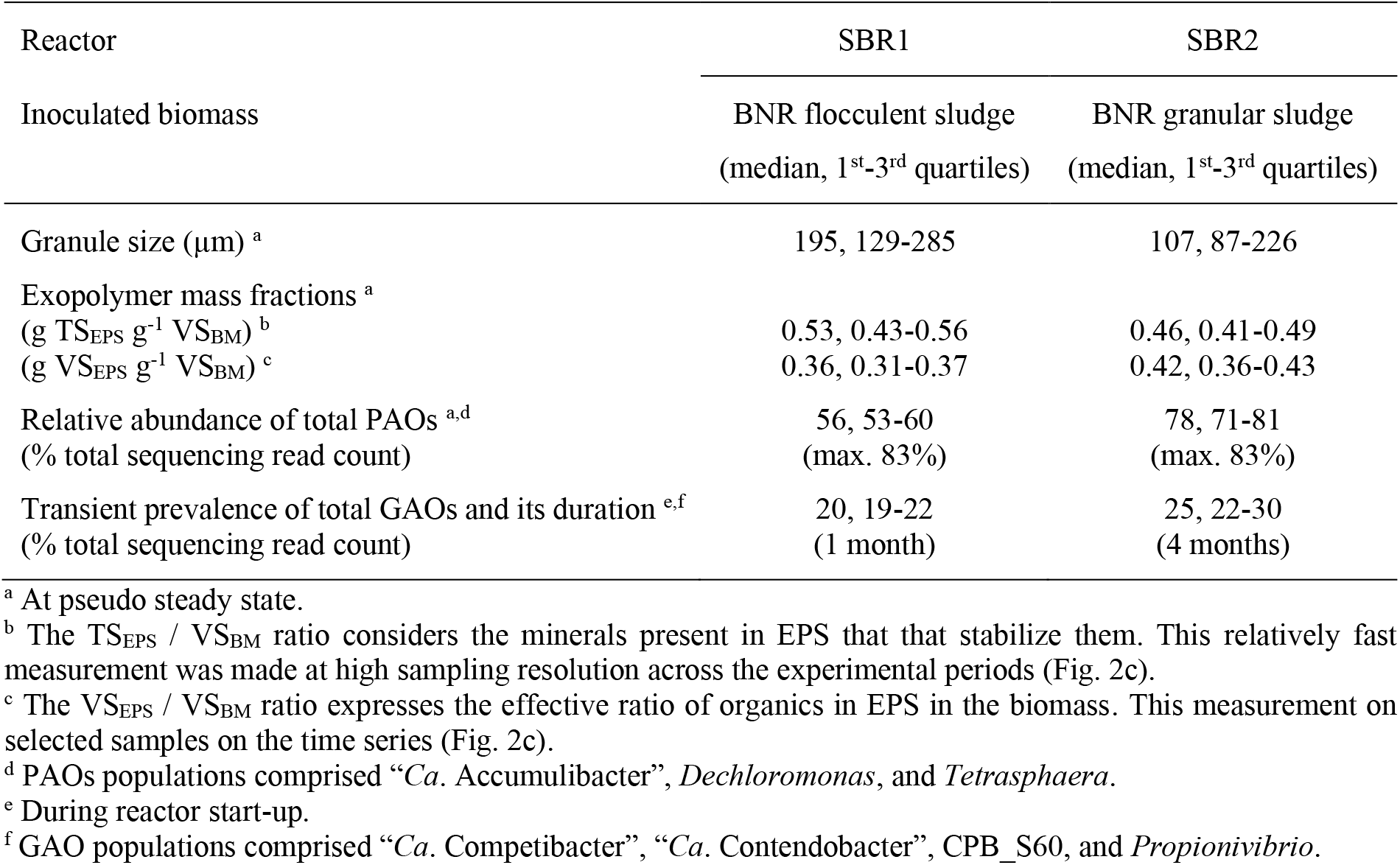
Characteristics of granules, exopolymers, and microorganisms formed under operation of the two bench-scale SBRs run under well-defined synthetic conditions. All values are given as distributions expressed by median and 1^st^-3^rd^ quartiles, over the start-up period (GAOs) and pseudo steady-state operation (granule size, exopolymer mass fractions, PAOs).

### 3.2 Exopolymers derived from PAO-enriched granules display a gel-forming behavior

The gel-forming property of the exopolymer extracts obtained from the granular biomasses of both SBRs was qualitatively validated by the preparation of sodium-based thin films and of typical gel beads morphology after chelation with divalent calcium cations (**Fig. 4a**).

**Figure 4.**
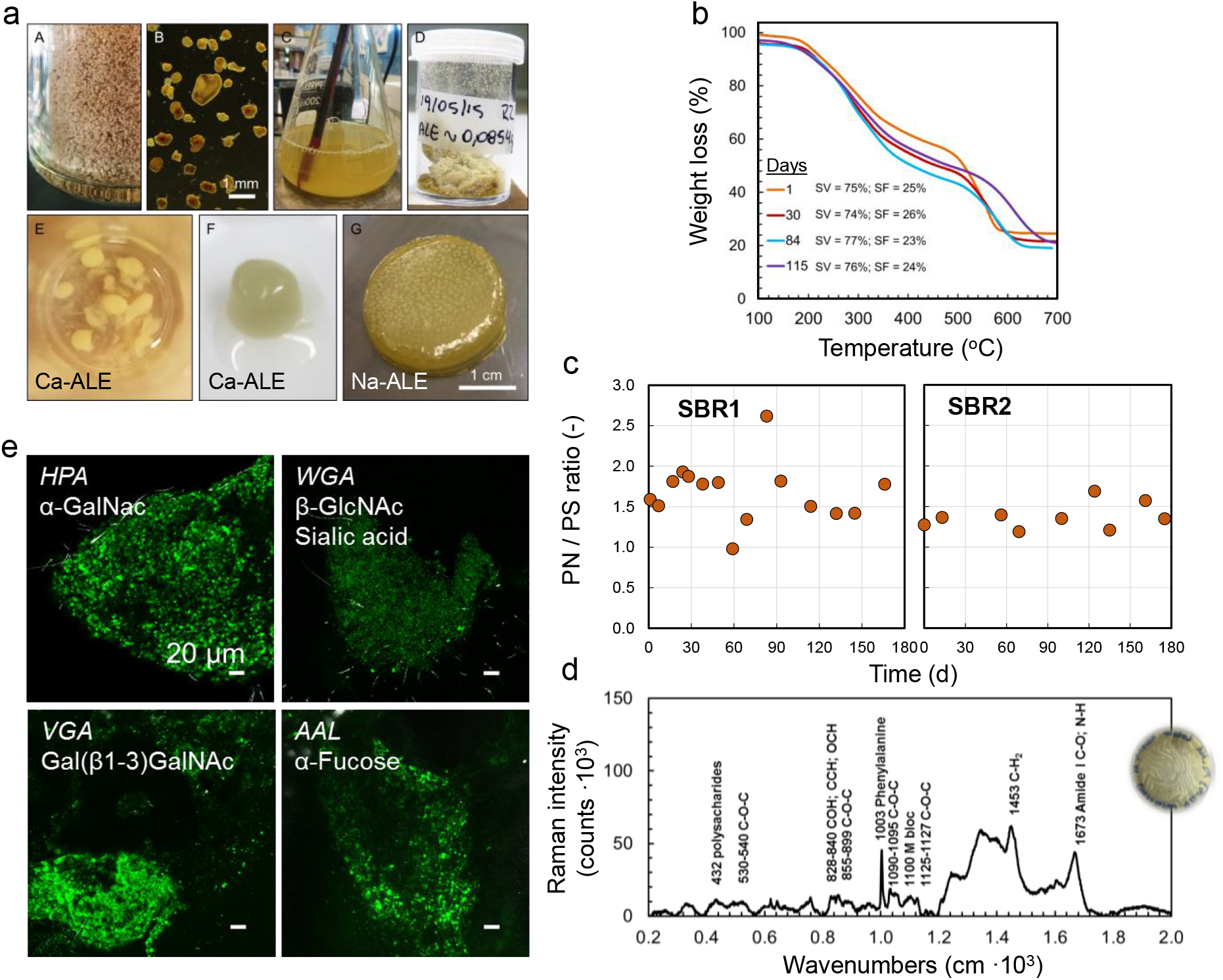
Material characteristics and chemical compositions of exopolymers recovered from granular sludge. Exopolymers were isolated from granules by physical-chemical extractions involving solubilization at pH ∼ 10 in a solution of sodium bicarbonate heated at 80°C, precipitation at pH 2.2, and lyophilization (**a** – *panels A-D*). The acidic exopolymers formed gel beads and gels when contacted with divalent calcium cations, and films with monovalent sodium cations (**a** – *panels E-G*). Thermogravimetry highlighted the thermal decomposition pattern of exopolymers from 100 to 700°C, with two main decomposition ramps at 200-350°C and 500-650°C (**b**). In the two SBRs, the exopolymers were composed of proteins, amino sugars, and neutral sugars, with a higher fraction of proteins, such as displayed by the protein-to-polysugar (PN/PS) ratio (**c**). The Raman spectrum of sodium-based films highlighted a complex chemical composition of exopolymers with peaks of glycosidic bonds of polysaccharides, of amino acids, of uronic acids and amides (**d**). Fluorescence lectin-binding analysis of granules cryosections detected abundant glycoconjugates containing amino sugars (detected with HPA, WGA and VGA lectins), sialic acids (WGA lectin), and fucose (AAL lectin) (**e**).

The thermogravimetric analysis displayed a thermal decomposition pattern of the exopolymer extracts from 100 to 700°C similar to alginate, with two main decomposition ramps from 200- 350°C and 500-650°C (**Fig. 4b**). The mass fraction of inorganics in the exopolymers increased in the first 60 days from 25 to 36% on samples collected from SBR1 but remained at low level and unchanged in 19 ± 1% in SBR2. Then, the ash fraction of the biomass was always higher than the ash fraction of exopolymers, either because of polyphosphate storage in the PAO cells or precipitation of minerals in granules. This led to higher mass fractions of exopolymers in the biomasses when measured as full VS ratios than as full TS ratios. On day 30, the difference between the mass fraction of exopolymers based on TS ratio and the one based on VS ratio was especially high. The VS-based mass fractions of exopolymers on day 30 and day 45 were equal (0.22 g VS_EPS_ g^-1^ VS_BM_), even though the TS-based mass fraction of exopolymers was lower on day 30 (120 g TS_EPS_ g^-1^ VS_BM_) than on day 42 (180 g TS_EPS_ g^-1^ VS_BM_).

### 3.3 The chemical compositions of EPS extracts fluctuate with population dynamics

#### Fluorescence lectin-binding analysis results

Fluorescence lectin-binding analysis (FLBA) of granules cryosections indicated the substantial presence of glycoconjugates containing amino sugars (α-GalNAc, β-GlcNAc, and Gal(β1- 3)GalNAc detected with the HPA, WGA, and VGA lectins, respectively), sialic acids (WGA lectin), fucose (AAL lectin) (**Fig. 4d**).

#### Raman and FT-IR spectroscopy results

Raman spectra of exopolymer sodium films displayed typical peaks of glycosidic bonds of polysaccharides, of uronic acids, of amino acids (phenylalanine) and amides (*i.e.*, presence of proteins) (**Fig. 4c**).

FT-IR spectra of the extracted EPS were analogous to the reference spectra of granular sludge and of exopolymers that have been reported earlier (Lin et al., 2010; Seviour et al., 2012; Lin et al., 2013). In both reactors, we identified the IR absorbance bands of typical chemical bonds of exopolymers at wavenumbers of 600, 812, 957, 972 (α-linkage between monomers), 1060 (pyranose rings), 1150, 1202, 1217, 1225 (O-acetyl ester), 1282, 1377 (CH bending), 1417, 1457 (CH2 bending), 1530 (C=C in pyranose ring), 1627 (asymmetric stretching of carboxylate O-C-O vibration / or amine peaks), 1720, 2862, 2917, 2950 (CH bond stretching), 3059, 3265 (hydroxyl group) cm^-1^. We noticed that the overall FT-IR spectra displayed considerable similarities over the whole operation time of the reactors (**Fig. 5**). Peaks around 1730 cm^-1^ and 1228 cm^-1^ are assigned to stretching vibration of C=O and C-O stretching vibration of the carboxyl group, respectively. Together with the results of the lectin staining presented in **Fig. 4a**, this can indicate the presence of sialic acid residues. The FT-IR spectra correlated with the operations of the two SBRs and the shifts in microbial selection. In both SBRs, the intensity of these bands at 1730 cm^-1^ and 1228 cm^-1^ increased when PAOs were predominant.

**Figure 5.**
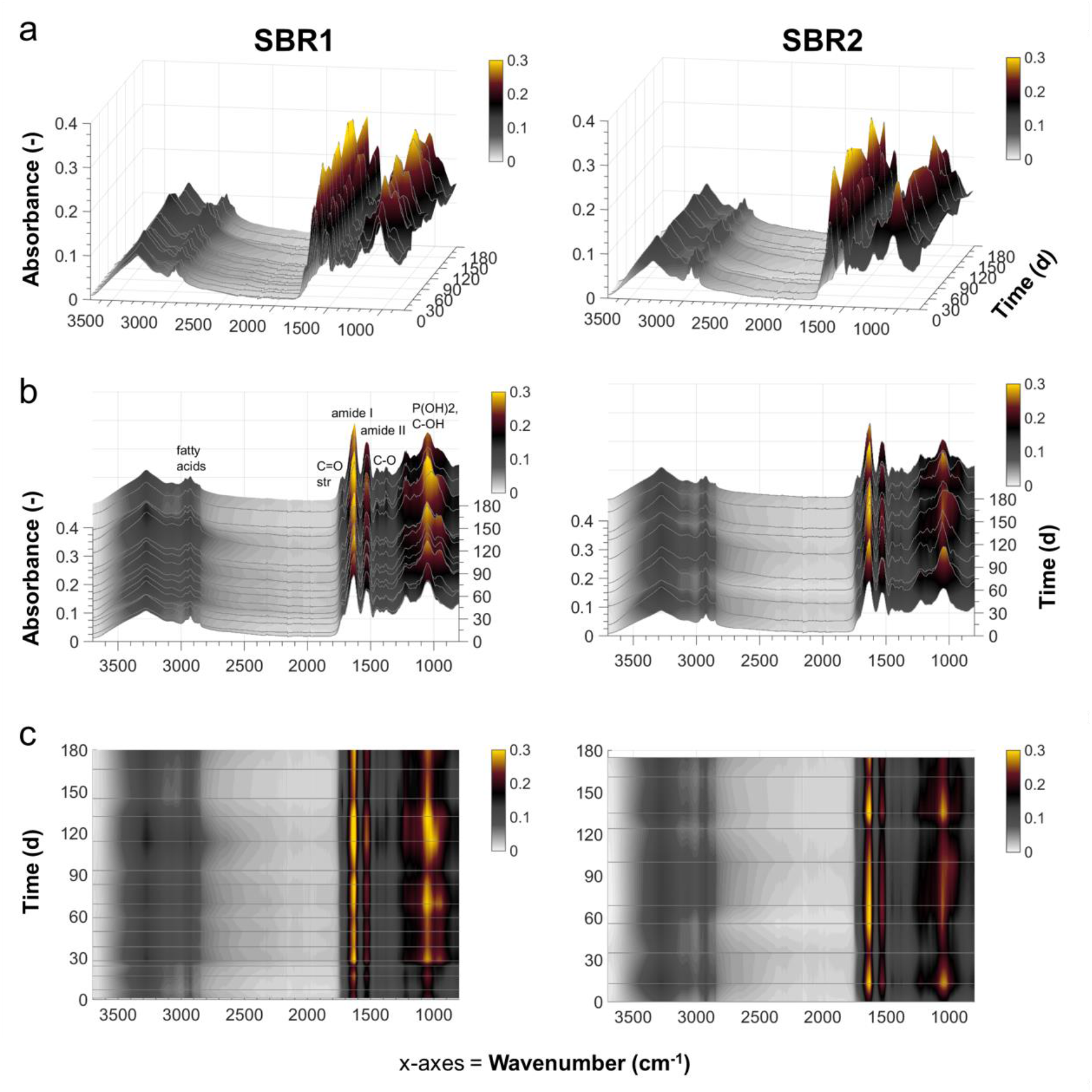
FT-IR spectral time series analyses (**a**- 3D view, **b**- front view, **c**- top view) of the overall chemical compositions of the exopolymers extracted from the granular biomasses cultivated over 180 days after inoculation with BNR activated sludge flocs in SBR1 (16 samples) and with BNR granular sludge in SBR2 (10 samples).

#### Profiling of sugars and aminosugars by HPAE-PAD chromatography

We qualitatively profiled the sugar and amino sugar monomers of the structural EPS and their dynamics along the reactors time series at high resolution with HPAE-PAD chromatography (**Fig. 6**). The EPS profiles displayed a complex assembly of monomers of glycerol, fucose, galactosamine, rhamnose, glucosamine, galactose, glucose, xylose, mannose, ribose, as well as the uronic acids galacturonic acid and glucuronic acid, in order of chromatographic retention times, among other unknown compounds. Interestingly, the sugars like fucose, rhamnose, galactose, xylose, mannose and ribose decreased over time in the EPS.

**Figure 6.**
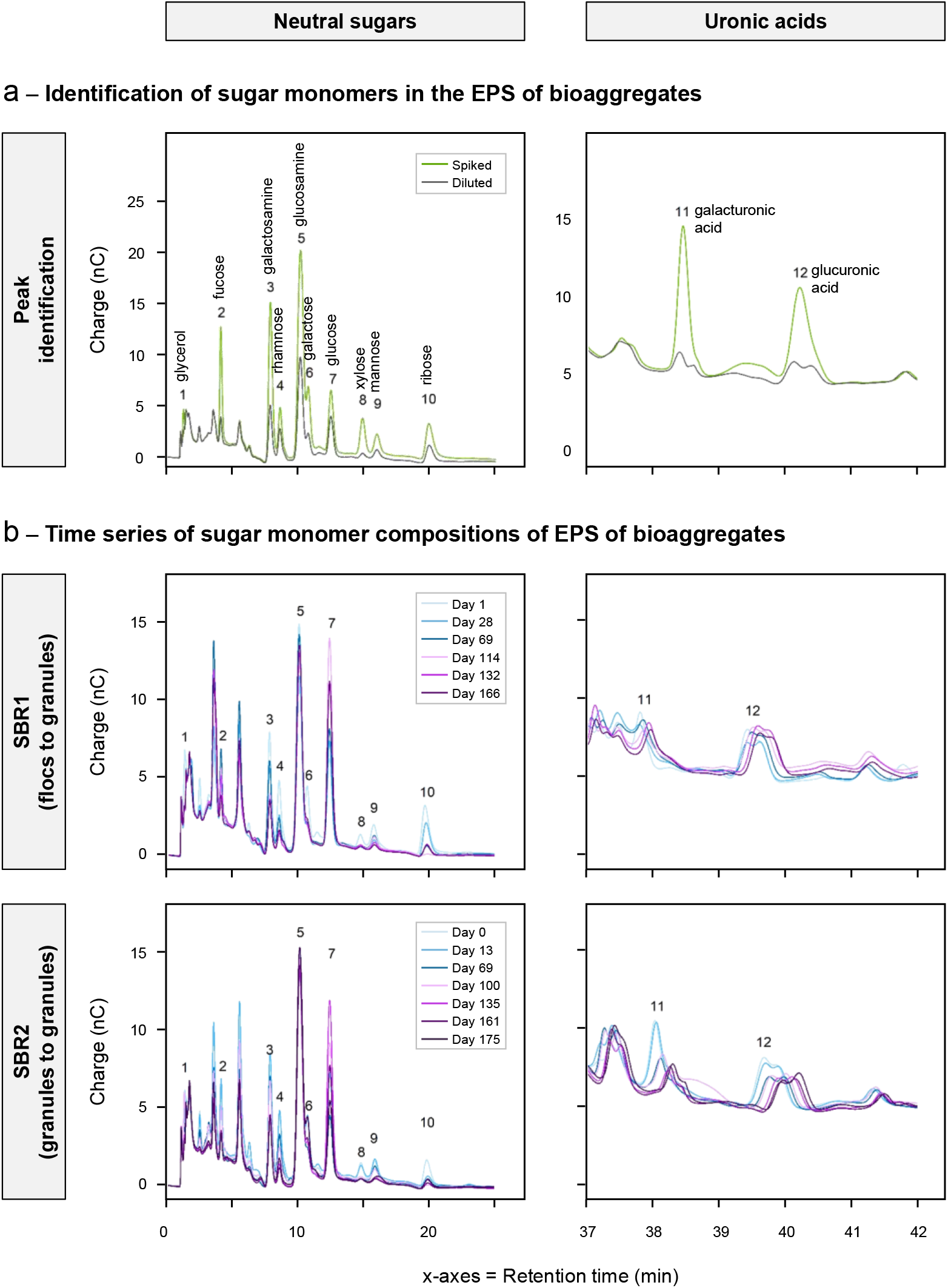
Qualitative analysis of the sugar monomers in the structural EPS by HPAEC-PAD chromatography. Detected monomers were (1) glycerol, (2) fucose, (3) galactosamine, (4) rhamnose, (5) glucosamine, (6) galactose, (7) glucose, (8) xylose, (9) mannose, (10) ribose. (11) galacturonic acid and (12) glucuronic acid. (**a**) Snapshots of the complexity of the sugar monomer compositions of the EPS (sample SBR1 day 1). The diluted analytes were spiked with internal standards for peak identification. (**b**) Temporal development of the sugar monomer compositions of the EPS of the bioaggregates (flocs, granules) across experimental time series of SBR 1 and SBR 2. The slight shift in peaks for the uronic acids (*right panels*) are inherent to the method used with elution with sodium acetate on batches of samples.

The composition profiles of microbial communities (**Fig. 3**) and chemical monomers (**Fig. 6**) were compared to detect the concomitant presence of microorganisms and EPS compounds. Monomers like glucose, galactose and galactosamine evolved along dynamics in “*Ca.* Accumulibacter” and “*Ca.* Competibacter”. In SBR1, glucosamine increased, and glucose decreased on day 114 in the EPS matrix when “*Ca.* Accumulibacter” became predominant. Galactose increased on day 132 when “*Ca.* Competibacter” re-entered the competition. In SBR 2, glucose and galactose decreased from day 135 onward in the EPS when “*Ca.* Accumulibacter” took over. The decrease of glucose and an increase of glucosamine in the structural EPS of EBPR granules can be related to the selection of “*Ca.* Accumulibacter”.

### 3.4 Genomic insights on the EPS metabolic potential of “*Ca*. Accumulibacter” and “*Ca*. Competibacter” lineages

Genetic signatures for EPS metabolisms in “*Ca.* Accumulibacter” and “*Ca.* Competibacter” lineages were compared to 46 other representative microbial genera present in activated and granular sludge microbiomes (Weissbrodt et al., 2014b; Winkler et al., 2018; Weissbrodt et al., 2020a) and to 9 model organisms known to produce exopolysaccharides for industrial biotechnological application (Schmid et al., 2015). A set of 20 metagenome-assembled genomes (MAGs) of “*Ca.* Accumulibacter” and 3 MAGs of “*Ca.* Competibacter” populations retrieved from public databases was compared to MAGs and genomes of the flanking and reference populations. More than 40 categories of functional genes coding for the metabolism of structural EPS were identified across these populations. Genetic annotations span broadly over the production and utilization of proteins, carbohydrates, cell wall and capsule, membrane transport, regulation and cell signaling across genomes. This highlights the complexity of the EPS biosynthetic pathways and their diversity across microbial populations of activated and granular sludges.

The **Fig. 7** provides a clustered heatmap of these functional gene categories potentially involved in the production and utilization of EPS. From this heatmap overview, “*Ca.* Accumulibacter” and “*Ca.* Competibacter” lineages form a relatively homogenous cluster of EPS genomic signatures (lineage clusters 4+5). The genomes of most of these PAO and GAO populations harbor a diversity of functional genes related to membrane surface and EPS components: (*i*) KDO2-lipid A biosynthesis, peptidoglycan biosynthesis, Type IV pilus, and Ton and Tol transport systems (genetic cluster 1; 17±6 instances); (*ii*) mycolic acid synthesis, cAMP signaling in bacteria, ABC transporter branched-chain amino acid (TC 3.A.1.4.1), and lacto-N-biose I and galacto-N-biose metabolic pathway (genetic cluster 2; 13±7 instances); (*iii*) sialic acid metabolism, colanic acid biosynthesis, linker unit-arabinogalactan synthesis, LOS core oligosaccharide biosynthesis, maltose and maltodextrin utilization (genetic cluster 4; 4±3 instances; this genetic cluster 4 is relatively conserved across all genomes of all microbial populations analyzed). The EPS-related genetic signatures of “*Ca.* Accumulibacter” and “*Ca.* Competibacter” populations clustered with well-known EPS producers like *Zoogloea*, *Thaurea* and *Azotobacter*, but also filamentous bacteria like *Sphaerotilus*, *Thiothrix* and *Leptothrix* genera present in wastewater environments.

**Figure 7.**
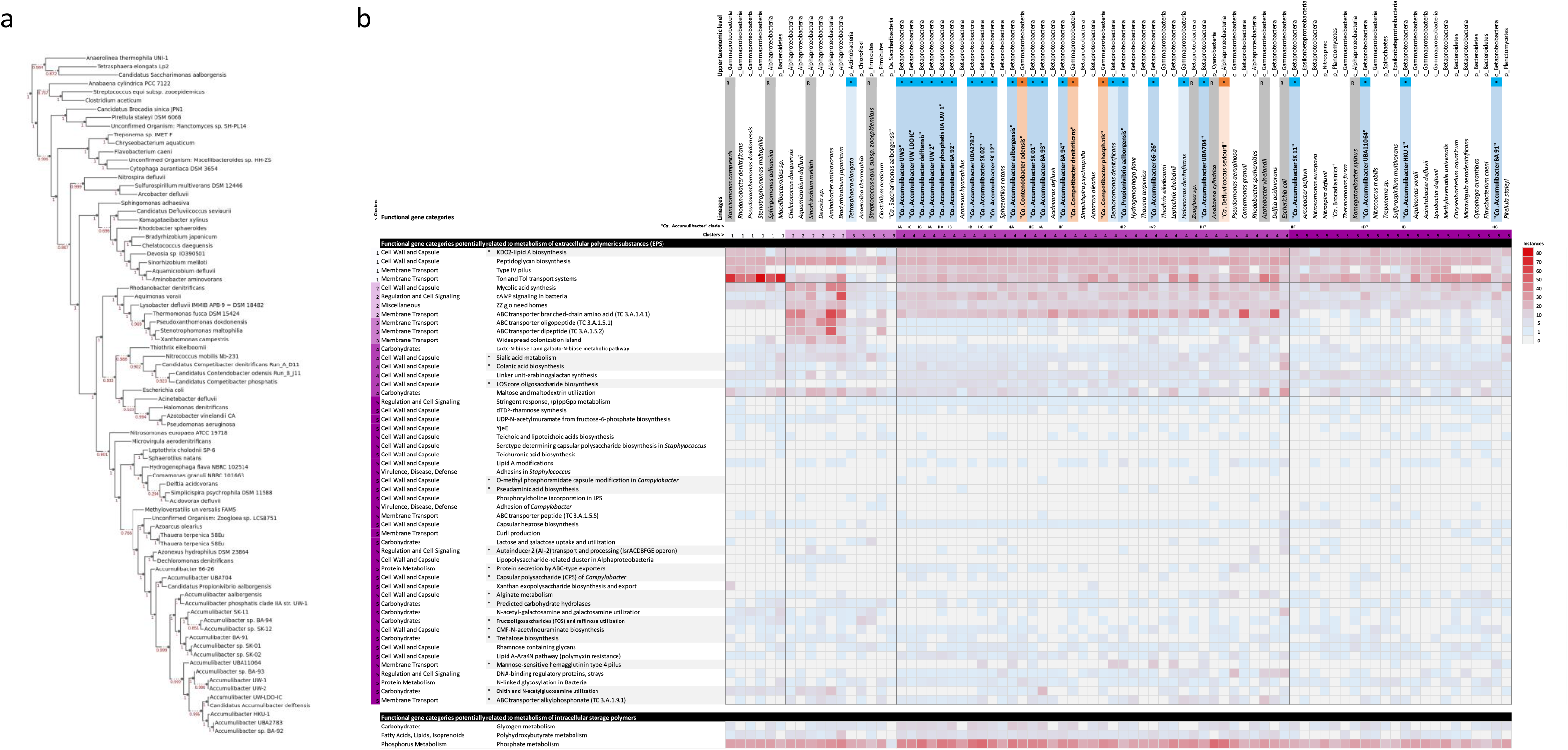
Identification of EPS genetic signatures in PAOs and GAOs. (**a**) Genome-based phylogenetic tree of representative lineages of “*Ca.* Accumulibacter” (highlighted in blue in b panel) and “*Ca.* Competibacter” (highlighted in orange) built with publicly available genomes according to Rubio-Rincon et al. (2019), among other populations of granular sludge selected from the ecosystem model of granular sludge (Weissbrodt et al., 2014b; Winkler et al., 2018; Weissbrodt et al., 2020a). (**b**) Clustered heatmap of functional gene categories potentially related to EPS metabolisms (production and utilization) in these microorganisms. Model organisms producing exopolysaccharides were used for comparison (highlighted in grey). Functions relating to the traditional metabolisms of glycogen, PHA and phosphorus in PAOs and GAOs are given as well.

The **Fig. 8** summarizes the main genetic categories present in the genomes of the lineage cluster 4 that comprises most populations of “*Ca.* Accumulibacter” and “*Ca.* Competibacter”. Besides signatures for peptidoglycan formation in cell membrane, these populations display biosynthetic potential for mycolic acid (long fatty acid), colanic acid, capsular heptoses, alginate, trehalose and rhamnose containing glycans (exopolysaccharides), sialic acid and CMP-N-acetylneuraminate (alpha-keto acid sugars), and N-linked glycosylation (binding of glycans to amino acids to form amino sugars or glycoproteins).

**Figure 8.**
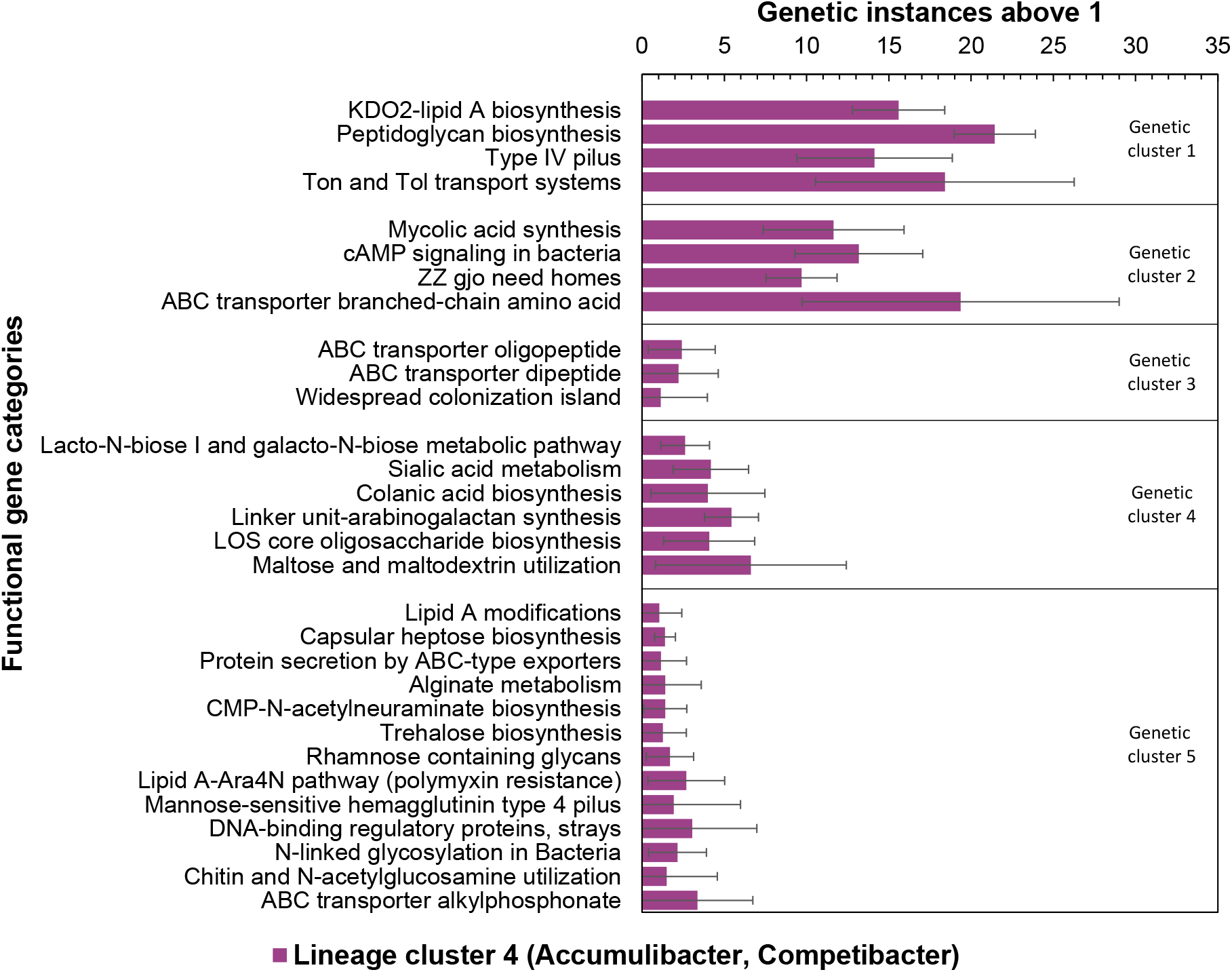
Functional gene categories possibly related to EPS metabolisms prevailing in the genomes of populations of “*Ca.* Accumulibacter” and “*Ca.* Competibacter” present in the lineage cluster 4 of **Fig. 7**.

## 4 Discussion

### 4.1 Microbial vectors of robust granulation

This study strengthened the consideration that slow-growing organisms forming compact microcolonies like the PAO “*Ca.* Accumulibacter” and GAO “*Ca.* Competibacter”, among other ones like nitrifiers and anammox organisms, are microbial vectors of a robust granulation (de Kreuk and van Loosdrecht, 2004; Weissbrodt et al., 2014a). Very compact and stable granules were produced under alternating anaerobic-aerobic conditions selecting for PAOs and GAOs, imposed by the operation of a strict anaerobic selector preventing the release of biodegradable COD in the aeration phase of the SBR.

The degree of enrichment of PAOs of up to as high as 83% of the amplicon sequencing read counts was triggered by the conditions applied for preferential selection in the lab-scale systems. Such high-grade enrichment may not be foreseen at full scale with municipal wastewater rich in particulate substrates. WWTPs connected to long and flat sewer systems where fermentation happens along wastewater transport can lead to substantial concentrations of VFAs in the influent, that can allow for PAO/GAO selection. Short and steep sewers result in a larger fraction of particulate substrates of up to 60% of the COD in the influent. Nonetheless, Ali et al. (2019) have shown that granules of a full-scale granular sludge can still be substantially enriched in PAOs, since particulate substrates mostly end up in the flanking floc fraction. Then, for all conditions, preferential selection for PAOs and GAOs will result from a duly operated anaerobic selector by managing the contact time between the influent wastewater and the biomass to enable hydrolysis, fermentation, and VFA uptake by these organisms prior to switching aeration or anoxic phases on. In granular sludge systems, this can occur via well-designed bed geometries and hydraulic conditions under slow up-flow feeding conditions, and/or by the integration of a subsequent anaerobic batch phase to promote the hydrolysis, fermentation, and uptake of VFAs by PAOs or GAOs (de Kreuk and van Loosdrecht, 2004; Weissbrodt et al., 2013b; Lochmatter and Holliger, 2014; Pronk et al., 2015b; Wagner et al., 2015; Derlon et al., 2016; Weissbrodt et al., 2017; Winkler et al., 2018; Toja Ortega et al., 2022; van Dijk et al., 2022).

Operation of SBR2 highlighted how BNR granules taken from a full-scale installation first disintegrated under the here-applied bench-scale operational conditions, prior to efficiently re- granulating under PAO-GAO selection. The cause for the initial disruption of full-scale granules remains puzzling. It can relate to different factors and combinations, *e.g.*, the: (*i*) high shear imposed by aeration in the up-flow bubble column, (*ii*) blocking of the nitrogen cycle by ATU and decay of microorganisms involved in nitrogen conversions, (*iii*) uncomplete penetration of the original bigger granules by the substrates, or (*iv*) more complex microbial and exopolymer compositions of the full-scale granules that get altered through the decay of several group of organisms outselected under operation under synthetic conditions. Mechanisms of granule formation, stability, and disruption have been conceptualized and modelled recently (Pronk et al., 2015a; van Dijk et al., 2022).

### 4.2 High exopolymer production under selection for slow-growing organisms

In addition to triggering granulation and EBPR, selection for “*Ca.* Accumulibacter” can propel a high production of exopolymers from used aqueous organic matter. The stepwise increase in the volumetric OLR from 0.3 to 2.0 g COD_Ac_ d^-1^ L_R_^-1^ and the adaptation of the COD:P ratio from 8 to 16 g CODAc g^-1^ P (*i.e.*, still in the safe range versus PAO-GAO competition (Schuler and Jenkins, 2003; Lopez-Vazquez et al., 2008; Weissbrodt et al., 2014a)) contributed to PAO selection and exopolymer production. Balancing the ratio of the most abundant divalent cations from 0.27 to 1 mol Ca^2+^ mol^-1^ Mg^2+^ sustained the accumulation of hydrogel-forming exopolymers, reaching a second plateau of up to 0.42 g VS_EPS_ g^-1^ VS_BM_ which is almost 2-3 times higher than reported for BNR granular sludge (Seviour et al., 2012; Lin et al., 2013; Pronk et al., 2017). In a control reactor operated under similar process conditions, similar EPS mass fraction have been obtained in a VFA-fed granular sludge (Schambeck et al., 2020).

Highest production of exopolymers correlated with an increasing predominance of PAOs and GAOs. This result was interestingly since these slow-growing chemoorganoheterotrophic organisms form compact microcolonies in activated sludge flocs and granules (Larsen et al., 2006; Weissbrodt et al., 2013a). The conjunction of microbial selection and organic loading was optimal to both generate compact granules and produce a substantial fraction of exopolymers in granular sludge. A high organic load has been evaluated as beneficial for the production of alginate-like exopolymers in granular sludge systems (Rollemberg et al., 2021).

Correlations were highlighted between the selection for PAOs and GAOs and the production of exopolymers. Different PAO and GAO lineages can be selected by the effect of wastewater compositions, and that can result in different granulation and exopolymer formation phenomena. The model PAO “*Ca.* Accumulibacter” and GAO “*Ca.* Competibacter” were selected by the VFA-based synthetic conditions and lower mesophilic temperatures (Lopez-Vazquez et al., 2009; Weissbrodt et al., 2013b; Weissbrodt et al., 2014a). In a previous SBR fed with acetate and operated at higher mesophilic temperature (35°C), granules have been enriched in other GAOs like *Defluviicoccus* and composed of capsular and acidic soluble EPS. They have displayed a different meso-scale architecture with a tangled tubular morphology (Pronk et al., 2017) rather than the conglomerates of microcolonies of “*Ca*. Accumulibacter” and “*Ca*. Competibacter” across the granules cross-sections (Weissbrodt et al., 2013a). More complex real wastewaters comprise a diverse mixture of dissolved and particulate organic and proteinaceous substrates (Kristiansen et al., 2013; Nguyen et al., 2015; Wagner et al., 2015) that are metabolized by a more diverse community of EBPR microorganisms (Nielsen et al., 2012). PAOs affiliating with *Tetrasphaera* and “*Ca.* Halomonas” are often predominant in activated sludge and granular sludge fed with real complex domestic wastewater (Nguyen et al., 2012; Nguyen et al., 2015; Layer et al., 2019). Like most of the microorganisms present in sludges, their exopolymer biosynthetic traits remain largely uncovered.

### 4.3 Chemical and monomeric variability of structural EPS in granular sludge biofilms

According to BCA and modified Dubois methods, the PN/PS ratio was constantly high in the EPS of BNR granules. It can be considered that the structural EPS of the granules was mainly proteins, and sugars that are linked to them. On top of protein and polysugar measurements, the high-resolution chemical profiling with Raman and FT-IR spectroscopies, and HPAEC-PAD chromatography highlighted the complex and chimeric chemical compositions of EPS extracts.

HPAEC-PAD was powerful to chemically profile the sugar momomers and to characterize their dynamics in the EPS products along SBR operations. The chemical and microbial time series revealed that the overall compositions of the EPS are linked to the operational conditions and microbial selection. This underlines the relationships between the microbial and chemical ecologies of granular sludge. Toward producing EPS products with defined compositions at full scale, temporal variabilities in wastewater characteristics and reactor operations need to be managed to master the chemical composition and physical properties of the recovered materials.

The increased intensity of bands associated with the carboxylic group in the FT-IR spectra, correlated with the increase in relative abundance of PAOs for both reactors. This chemical functional group should be further investigated in granular sludge. The presence of sialic acid residues in granular sludge and other wastewater biomasses has been previously reported and described (Weissbrodt et al., 2013a; de Graaff et al., 2019; Kleikamp et al., 2020; Tomás-Martínez et al., 2021). These nonulosonic acids linked to a carbohydrate chain can exert a protective role against biodegradation. From the chromatographic measurements, we related the EPS amino sugar glucosamine to the abundance of “*Ca.* Accumulibacter”. Most of sugar residues dropped after inoculation with full-scale BNR flocs or granules and transient pre-selection of “*Ca.* Competibacter”. N-acetylglucosamine has been detected earlier by FLBA- CLSM in cross-sections of granules enriched in “*Ca.* Accumulibacter” (Weissbrodt et al., 2013a). These amino sugars can be considered as preferential functional and structural EPS used by “*Ca.* Accumulibacter” to bioaggregate, while “*Ca.* Competibacter” uses glucose and galactose. EPS biosynthetic signatures are specific to microbial populations, such as shown for exopolysaccharides (Schmid et al., 2015). Individual lineages may further synthesize different EPS depending on growth and environmental conditions they encounter. Overall, the EPS-related genetic information screened from the public MAGs of “*Ca.* Accumulibacter”, “*Ca.* Competibacter” and other representative genera of wastewater ecosystems clustered relatively closely at functional genetic groups. More detailed, targeted analyses of EPS biosynthetic gene clusters can reveal the fine-scale genetic differentiations, following the recent developments by Dueholm et al. (2023).

In addition, as shown by our measurements, proteins occupy a large fraction of the EPS mass, and substantial work will be needed to unravel their compositions and metabolisms. EPS are not solely related to exopolysaccharides, with predominant proteins and glycoproteins making the analytical workflow even more complex (Weissbrodt et al., 2013a; de Graaff et al., 2019; Boleij et al., 2020; Felz et al., 2020). Analytical challenges remain across extraction, purification, separation, and detection workflows to identify and sequence EPS with high quantitative accuracy (Felz et al., 2016; Seviour et al., 2019). High-throughput chromatographic analyses of carbohydrate and amino sugar monomers (Felz et al., 2019) will help characterize and manage the compositions of EPS products. The relationships between the (varying) chemical compositions and physical properties of the polymers deserve a closer look. Exopolymer compositions are as complex as microbial communities in biological wastewater treatment (Weissbrodt et al., 2014b; Winkler et al., 2018).

### 4.4 Genomic and functional determinants of EPS biosynthesis in complex communities

Beyond the correlative analysis of microbial and chemical ecologies of granules, we addressed the genomic and biosynthetic signatures of the model PAO “*Ca.* Accumulibacter” and GAO “*Ca.* Competibacter” for EPS formation, in order to predict their functional involvement among other activated sludge and granular sludge populations. We examined representative genomes of “*Ca.* Accumulibacter” and “*Ca.* Competibacter” available in public repositories to analyze their genetic functions related to EPS.

EPS formation involves a diversity of metabolic pathways, intermediates, and products from carbohydrates to amino sugars, glycoproteins and proteins. Most of genetic investigations of EPS biosynthesis have been performed on exopolysaccharides of health and plant pathogens and microorganisms relevant for industrial biotechnology (Schmid et al., 2015; An et al., 2016). The rest mostly remains a *terra incognita*. At exopolysaccharides level, the genomes analyzed from “*Ca.* Accusmulibacter” and “*Ca.* Competibacter” and their flanking populations highlighted that EPS metabolisms rely on a diversity of functional categories harbored by these populations. A closer look to functional genes indicate that the three main biosynthetic pathways described by Schmid et al. (2015). The *Wzx/Wzy-dependent pathway* relate to the metabolisms of UDP-glucose, UDP-glucuronic acid, dTDP-L-Rhamnose, o-acetate, phosphate, and monomers of colanic acid (Glc, Fuc, GlcA, Gal) and xanthan (Glc, Man, GluA) involving the GTs, Wzx, Wzy, PCP, Wzz, OPX proteins. The *ABC transporter-dependent pathway* links to UDP-glucose, UDP-glucuronic acid, dTDP-L-rhamnose, kdo-linker, phosphate, involving the GTs, ABC transporter, PCP, OPX proteins. The *synthase-dependent pathway* relate to UDP-glucose and monomers of alginate (GulA, ManA), cellulose (Glc) and hyaluronic acid (GlcA, GlcNAc), involving synthases, c-di-GMP, TPR, and beta-Barrel proteins.

The task of elucidating EPS biosynthetic pathways is not as simple as for the investigation of, *e.g.*, catabolic pathways within the nitrogen or phosphorus cycle of wastewater microorganisms (Gao et al., 2019; Rubio-Rincon et al., 2019). The lack of a comprehensive database of highly annotated EPS genes in wastewater microorganisms has been reported as one main limitation (Albertsen et al., 2013b; Barr et al., 2015; Dueholm et al., 2023). Databases of high-quality MAGs from sludge metagenomes (Singleton et al., 2021) developed using genome-centric metagenomics (Albertsen et al., 2013a) now allow for elucidating at higher resolution the genetic potential of wastewater microorganisms for EPS biosynthesis, like exopolysaccharides (Dueholm et al., 2023). Still only 11% of the 1000 HQ MAGs examined by the authors encoded homologs to known exopolysaccharide gene clusters, highlighting the pressing need to further resolve this EPS and their genetic dark matter. Beyond the genetic potential, experiments should be designed to examine expressed EPS functions. Instead of addressing the broad diversity of EPS metabolic signatures across the biomass, one rational approach can target a set of specific EPS compounds. This was recently started with the analysis of nonulosonic acids in granular sludge by mass spectrometry and metaproteomics (Kleikamp et al., 2020; Kleikamp et al., 2021; Tomás-Martínez et al., 2021). Combination with ^13^C substrate labelling experiments help examine their metabolic turnover (Tomás-Martínez et al., 2023b). Such advanced analytical workflow will drive a stepwise elucidation of the complex metabolisms of EPS in PAOs, GAOs, activated sludge populations, and microbial communities in general.

### 4.5 Toward an ecological engineering of exopolymer production in mixed cultures

Exopolymers obtained in granules formed under selection for PAOs and GAOs displayed interesting material and gelling properties. Structural EPS and other valuable materials extracted from granular biomass can find industrial application as coatings in paper, concrete, fiber manufacturing and as nanocomposite flame-retardant material when combined with nano- clay discs (Lin et al., 2015; Zlopasa et al., 2015; Kim et al., 2020; Espíndola et al., 2021). These waste-based biomaterials can function as water-resistant film to protect hydrophilic surfaces such as paper, extend the lifespan of concrete, and form clothing fibers after purification. Future developments of mixed-culture processes for the production of such hi-tech exopolymers will benefit from the implementation of ecological engineering principles (Kleerebezem and van Loosdrecht, 2007; Angenent and Kleerebezem, 2011; Lawson et al., 2019), by controlling the selection of microorganisms, their metabolisms, and their EPS products.

## 5 Conclusions

We provided insights into the relationships between microbial selection, bioaggregation, as well as EPS production and compositions in BNR granular sludge. Collectively, our results led to the following main findings:

- Granulation was promoted and stabilized by the selection of slow-growing organisms like the model PAO “*Ca.* Accumulibacter” and GAO “*Ca.* Competibacter” in anaerobic-aerobic SBRs. A strict anaerobic selector was achieved here by complementing the up-flow feeding of wastewater across the biomass bed by a subsequent anaerobic batch phase.
- Substantial production of EPS was obtained under these conditions up to a mass fraction as high as 0.40 g VS_EPS_ g^-1^ VS_BM_ (about 4 to 6-fold higher than in activated sludge flocs).
- A resistant matrix of structural EPS was built under predominance of PAOs, which was resistant to pH shocks. The calcium-mediated gelation of EPS extracts validated their attractive biomaterial property.
- Chemical composition of EPS is complex, as a combination of proteins, glycoproteins, amino sugars, and sugars, and varied in function of microbial selection and process conditions. Managing seasonal variations in wastewater characteristics and process operation will be needed to direct the compositions of the EPS products.
- Genetic signatures of PAOs and GAOs for the metabolisms of EPS were annotated from publicly available genomes among of other representative organisms of granular sludge. Future investigations of functional expression and metabolic regulations will help identify, and potentially control, the main EPS biosynthetic pathways involved in mixed cultures.

Our findings can be utilized to design experiments in order to better capture the metabolisms of EPS in mixed cultures. It provides scientific insights to current incentives that expand granular sludge systems as biorefineries to valorize exopolymers on top of an intensified environmental service, transforming sludge disposal into a resource of valuable hi-tech biomaterials.

## Acknowledgements

This study was financed by the Swiss National Science Foundation, grant no. 151977 (David Weissbrodt) and supported by internal funds at Delft University of Technology and Aalborg University. Lorena Guimarães was supported by CAPES and CNPq doctoral grants in Brazil and for a doctoral research stay in Delft and Aalborg. Nina Gubser was recipient of the IDEA League master grant at ETH Zürich in Switzerland for her MSc thesis conducted in Delft and Aalborg. Udo van Dongen, Yang Song and Dirk Geerts in Delft as well as Jane Ildal, Jannie Munk Kristensen and Søren Karst in Aalborg are acknowledged for their support on bioprocess setup and molecular methods, respectively. We thank RWZI Harnashpolder and Royal HaskoningDHV in The Netherlands for providing the activated sludge and granular sludge, respectively, used to seed the bioreactors.

## Author contributions

DGW, NRG and LBG designed the study, processed the data, and extracted the scientific findings, with inputs of MCMvL, PHN and RHRdC. LBG and NRG operated, sampled, and analyzed the bioreactors, supported by DGW and MP. LGB, NRG and DGW performed the wet-lab and dry-lab analyses of molecular microbial ecology analyses in collaboration with by MA, MKDD and PHN. LBG and NRG performed the extractions and characterizations of EPS together with YL, JZ and SF. TRN performed the FLBA-CLSM screening of glycoconjugates. DGW made the bioinformatics computations to examine EPS signatures in genomes, with inputs of MA, MKDD, STM and PHN. LBG and DGW wrote the manuscript with contribution, edits, and critical feedback by the co-authors.

## Preprint

The manuscript is deposited as pre-print in *bioRxiv*.

